# Host Enzymes Heparanase and Cathepsin L Promote Herpes Simplex Virus-2 Release from Cells

**DOI:** 10.1101/364562

**Authors:** James Hopkins, Tejabhiram Yadavalli, Alex M. Agelidis, Deepak Shukla

## Abstract

Herpes simplex virus-2 (HSV-2) can productively infect many different cell types of human and non-human origin. Here we demonstrate interconnected roles for two host enzymes, heparanase (HPSE) and cathepsin L in HSV-2 release from cells. In vaginal epithelial cells and other cell lines tested, HSV-2 causes heparan sulfate shedding and upregulation in HPSE levels during the productive phase of infection. We also noted increased levels of cathepsin L and show that regulation of HPSE by cathepsin L via cleavage of HPSE proenzyme is important for infection. Furthermore, inhibition of HPSE by a specific inhibitor, OGT 2115, dramatically reduces HSV-2 release from vaginal epithelial cells. Likewise, we show evidence that the inhibition of cathepsin L is detrimental to the infection. The HPSE increase after infection is mediated by an increased NF-κB nuclear localization and a resultant activation of HPSE transcription. Together these mechanisms contribute to the removal of heparan sulfate from the cell surface, and thus facilitate virus release from cells.

**Importance:** Genital infections by HSV-2 represent one of the most common sexually transmitted viral infections. The virus causes painful lesions, and sores around the genitals or rectum. Intermittent release of the virus from infected tissues during sexual activities is the most common cause of transmission. At the molecular level, cell surface heparan sulfate (HS) is known to provide attachment sites for HSV-2. While the removal of HS during HSV-1 release has been shown, not much is known about the host factors and their regulators that contribute to HSV-2 release from natural target cell types. Here we suggest a role for the host enzyme heparanase in HSV-2 release. Our work reveals that in addition to the regulation of transcription by NF-κB, HPSE is also regulated post-translationally by cathepsin L and that inhibition of heparanase activity directly affects HSV-2 release. We provide unique insights into the host mechanisms controlling HSV-2 egress and spread.

## Introduction

Genital herpes is one of the most common, persistent and highly infectious sexually transmitted disease caused by herpes simplex virus type-2 (HSV-2) and in many emerging first-time cases, by herpes simplex virus type-1 (HSV-1)(1-4). Primarily, the sites of infection include the vulva and the vagina, with some cases involving the cervix and perianal region in women and typically on the glans or the shaft of the penis in heterosexual men, whereas anal infection has also been reported with homosexual men (5-7). Primary and recurrent genital herpes infections result in lesions and inflammation around the genital area which are painful and cause distress (4). While there is no vaccination or cure against HSV-2, resistance against current therapies, such as Acyclovir, have been reported (8). Furthermore, these therapies are more than a decade old and work on a single aspect of the viral life cycle, viral DNA replication. Novel therapeutic interventions that target different stages of viral infection including viral entry, viral protein translation and viral egress need to be addressed to successfully curb this distressing disease. One method to generate novel antiviral drugs that target these viral pathways is to understand host factors that help facilitate viral lifecycle. In this manuscript we focus on the host enzyme heparanase (HPSE) and its regulators that help facilitate egress of the HSV-2 virions.

Human HPSE is an endoglycosidase with the unique distinction of being the only enzyme capable of degrading heparan sulfate (HS)(9-11), an evolutionarily conserved glycosaminoglycan that is present ubiquitously at the cell surface. HPSE is initially translated as a pre-proenzyme. Cleavage of a signal sequence by a signal peptidase leaves an inactive 65 kDa proHPSE, which undergoes further processing in the lysosomal compartment (10, 12). Proteolytic removal of an N-terminal 8 kDa linker by a lysosomal cysteine endopeptidase, cathepsin L, cleaves the C-terminal 50 kDa subunit, which remain associated as a non-covalent heterodimer in active HPSE(13, 14). Active HPSE is responsible for the degradation of cell surface HS, that is found covalently attached to a small set of extracellular matrix and plasma membrane proteins forming heparan sulfate proteoglycans (HSPG) (15, 16). Clearance of HS via HPSE modulates cell division and differentiation, tissue morphogenesis and architecture, and organismal physiology (16). HSV-2 encodes for two envelop glycoproteins, gB and gC, which bind HS at the cell surface and initiate viral entry (17-19). We first reported that host-encoded HPSE is upregulated and required for the release of viral progeny from parent cells after HSV-1 infection and subsequently similar findings were reported for porcine reproductive and respiratory syndrome virus (PRRSV) infection(20, 21).

The premise of this manuscript is to understand the role of HPSE in the egress of HSV-2 virus from its natural target cells. In this study, we show that HPSE is upregulated by the virus upon infection and serves to aid in viral egress by preventing the newly released viral progeny from reattaching to cell surface HS. We also study transcriptional and post-translational regulators of HPSE and for the first time implicate cathepsin L in HSV release. We demonstrate that inhibition of HPSE and cathepsin L via commercially available inhibitors negatively impacts viral egress.

## Results

### Loss of cell surface HS during infection

To understand how HSV-2 infection modulates cell surface HS levels, we infected a natural target cell type, human vaginal epithelial cells (VK2), with HSV-2 with a multiplicity of infection (MOI) of 1 for a period of 48 hours. We observed that while mock infected cells consistently showed high amounts of cell surface HS, most HS was cleared in HSV-2 infected cells by 24 hours post infection (hpi) (Figure 1A, 1B). We also observed that there was a progressive loss of HS on the cell surface with time during infection using flow cytometric analysis (Figure 1C, 1D).

**Figure 1.**
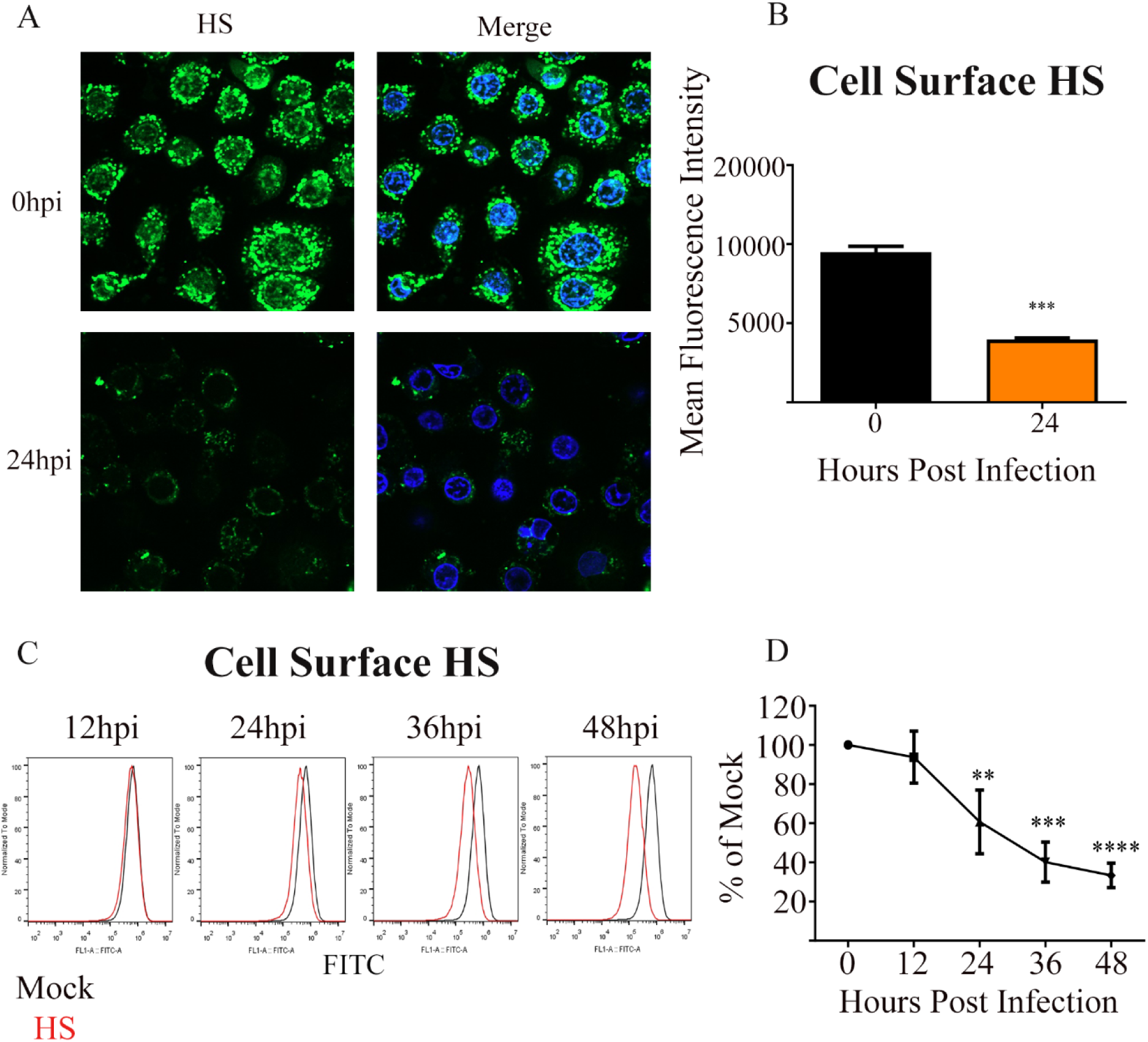
Loss of cell surface HS during infection. **A**. Representative immunofluorescent images of HS stain. HSV-2 333 was used to infect cells at 1MOI for 24h. Upper left is HS stain only in uninfected sample, upper right is hoescht and HS stain merged for uninfected, lower left is HS stain only for infected sample at 24hpi, lower right is hoescht and HS stain merged for infected sample at 24hpi. **B**. Quantification of HS cell surface expression. **C**. Representative flow cytometry histogram showing change in cell surface HS expression with red representing infected samples and black showing uninfected control. **D**. Quantification of cell surface HS flow cytometry experiments. Asterisks denote a significant difference as determined by Student’s *t*-test; ^∗^*P*<0.05, ^∗∗^*P*<0.01, ^∗∗∗^*P*<0.001, ^∗∗∗∗^P<0.0001.

### HPSE is upregulated after HSV-2 infection

Given that clearance of HS correlated with duration of HSV-2 infection, we hypothesized an upregulation of HPSE expression. HPSE is the only mammalian enzyme known to cleave HS. In order to test our hypothesis, we analyzed HPSE promoter activity at 12, 24, 36, and 48 hpi using a luciferase reporter assay. As expected, we observed a significant increase in the HPSE promoter activity during the duration of infection (Figure 2A). To understand this further we then looked at HPSE mRNA transcript levels after HSV-2 infection. We observed by quantitative real time-PCR (qRT-PCR) analysis that HPSE mRNA was significantly elevated at 12, 24 and 36 hpi (Figure 2B). Since HPSE expression was clearly upregulated inside the cells, we decided to look at HPSE translocation to the surface, which is the primary site for HS removal (16). Complementary to our HPSE mRNA results, after infection at an MOI of 1 there was a significant increase in cell surface HPSE protein levels that was observed first by immunofluorescence microscopy that showed a large and significant increase at 24 hpi (Figure 2C, 2D). This increase in cell surface levels of HPSE was subsequently verified by flow cytometry (Fig. 2E, 2F). Taken together, our results confirmed an upregulation in HPSE levels upon infection.

**Figure 2.**
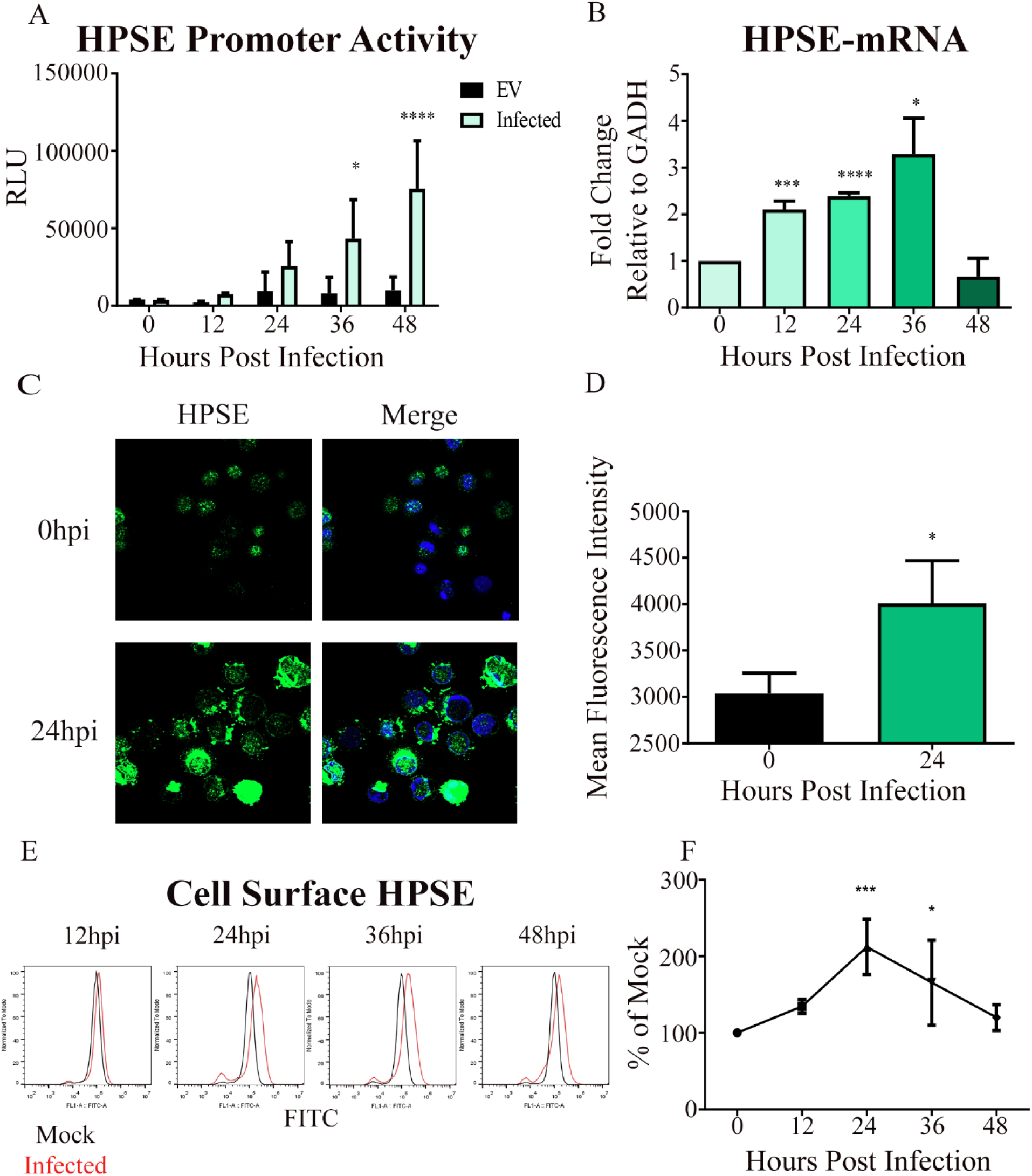
HPSE is upregulated after HSV-2 infection. **A**. Increase in promoter activity of HPSE gene upon infection in HCE cells. HSV-2 333 was used to infect cells at a MOI of 1 for 12, 24, 36 and 48h. Shown is the average fold increase over uninfected control. Experimental values are normalized to those obtained with pGL3 as a control for transfection efficiency. **B**. Increase in HPSE mRNA levels, shown is the average fold increase over uninfected control. **C**. Representative immunofluorescence microscopy images of cell surface HPSE stain. HSV-2 333 was used to infect cells at a MOI of 1 for 24h. Upper left is HPSE stain only in uninfected sample, upper right is hoescht and HPSE stain merged for uninfected, lower left is HPSE stain only for infected sample at 24hpi, lower right is hoescht and HPSE stain merged for infected sample at 24hpi. **D**. Quantification of HPSE cell surface expression from multiple immunofluorescent images. **E**. Representative flow cytometry histogram showing change in cell surface HPSE expression with red representing infected samples and black showing uninfected control. **F**. Quantification of cell surface HPSE flow cytometry experiments. Asterisks denote a significant difference as determined by Student’s *t*-test; ^∗^*P*<0.05, ^∗∗^*P*<0.01, ^∗∗∗^*P*<0.001, ^∗∗∗∗^P<0.0001.

### Mechanism of HPSE upregulation and activation upon infection

To understand the mechanism of HPSE regulation during HSV-2 infection, we studied transcriptional and post-translational regulators of HPSE. It is reported that during HSV-1 infection nuclear factor NF-κB (p65) activation and translocation to the nucleus could transcriptionally increase HPSE expression (22, 23). To understand if HSV-2 used a similar mode of action, we analyzed p65 activity during HSV-2 infection. As expected, there was a consistent increase in p65 mRNA expression in HSV-2 infected cells through 36 hpi (Figure 3A). We also observed the nuclear translocation of p65 (Figure 3B) at 24 hpi using immunofluorescence microscopy. These results corroborated our western blot data, which showed significantly increased p65 in the nuclear fraction with concurrent decrease in the cytoplasmic fraction after HSV-2 infection (Figure 3D). Next we wanted to understand if the inhibition of p65 activation and nuclear localization would affect HPSE promoter activity. In this regard, we overexpressed a plasmid encoding mutant IkBa (S32A/S36A) in VK2 cells. This mutant IkBa is incapable of being phosphorylated and degraded and as a result, it acts as a dominant-negative protein that inhibits NF-κB activation and nuclear translocation (24). We did observe that expression of the IkBa mutant lead to a decrease in HPSE promoter activity (Figure 3C). Taken together our findings suggest that virus-induced activation of NF-κB regulates HPSE expression.

**Figure 3.**
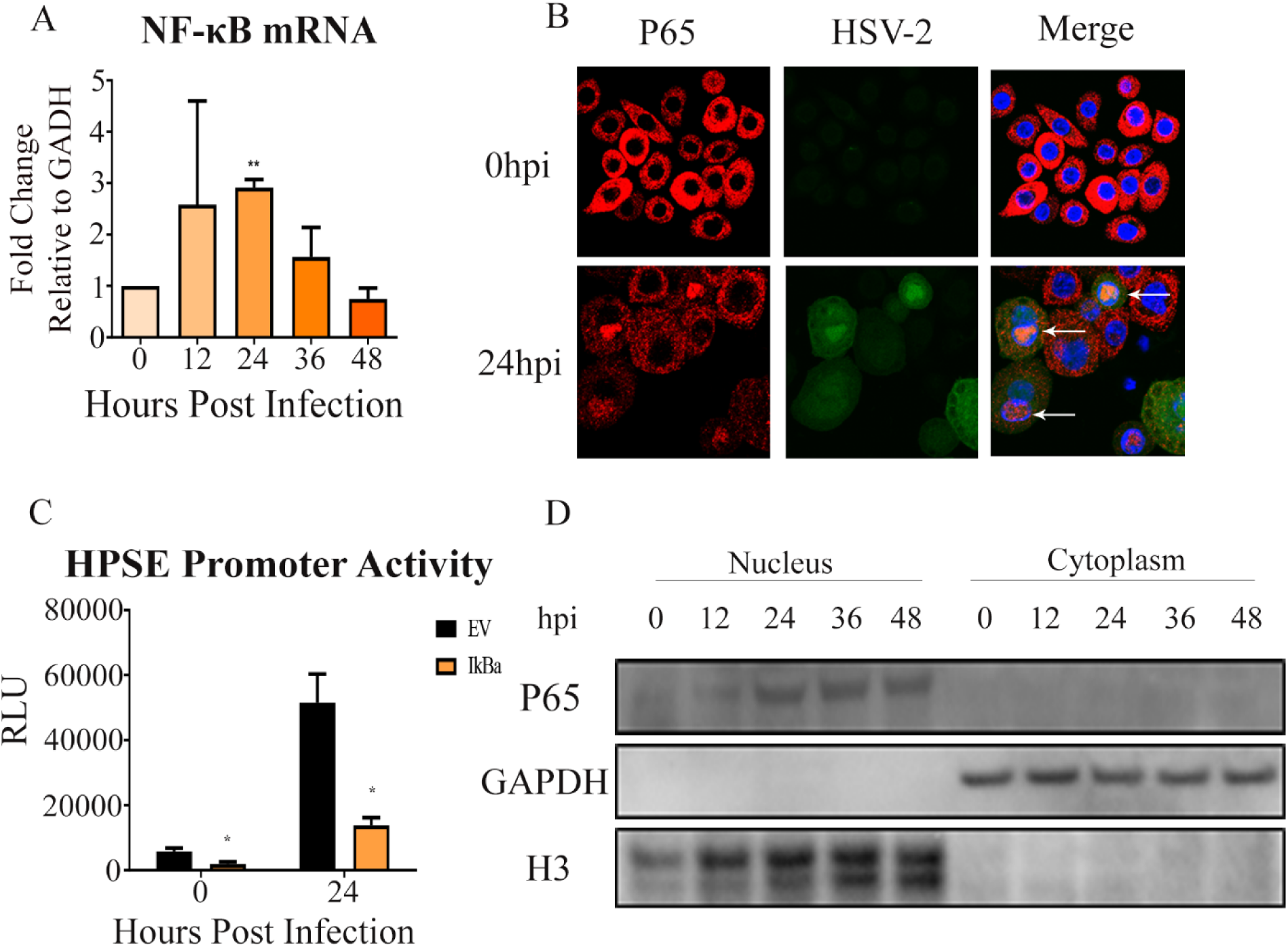
Nf-κB as mechanism for HPSE upregulation. **A**. Increase in Nf-κB P65 mRNA levels, shown is the average fold increase over uninfected control. **B**. Representative immunofluorescence microscopy images of nuclear translocation of P65 upon HSV-2 infection with HSV-2 333 GFP. With Nf-κB P65 labeled red, HSV-2 green, and the nucleus blue. **C**. Inhibition of NF-κB activation results in decreased HPSE promoter activity. VK2 cells were transfected with mutant IkBa incapable of degradation (S32A/S36A), thereby specifically inhibiting NF-κB activation and nuclear translocation. 24h after transfection cells were infected with HSV-2 333 for 24 hrs at a MOI of 1. Cell lysates were isolated, and luciferase assay was performed. Results shown are normalized to empty pGL3 vector as a control for transfection efficiency. **D**. Representative western blot of nuclear translocation of P65 upon HSV-2 infection with HSV-2 333. Asterisks denote a significant difference as determined by Student’s *t*-test; ^∗^*P*<0.05, ^∗∗^*P*<0.01.

As described by others in the field, cathepsin L is the only known post-translational activator of HPSE in mammalian cells(13, 25). Given that HPSE is upregulated upon HSV-2 infection, we wanted to assess whether its lysosomal activator, cathepsin L, also increased with HSV-2 infection. As hypothesized, cathepsin L mRNA levels were upregulated at 12 hpi when compared to mock and continued to increase at 24 and 36 hpi (Figure 4A). We observed a similar trend in cathepsin L protein levels where the maximum protein expression was seen at 48 hpi by western blot analysis (Figure 4B, 4C). These important findings were confirmed using immunofluorescence microscopy (figure 4D, 4E) and flow cytometry analysis (Figure 4F, 4G). Collectively, our data suggests a connection between HSV-2 infection and HPSE/cathepsin L upregulation.

**Figure 4.**
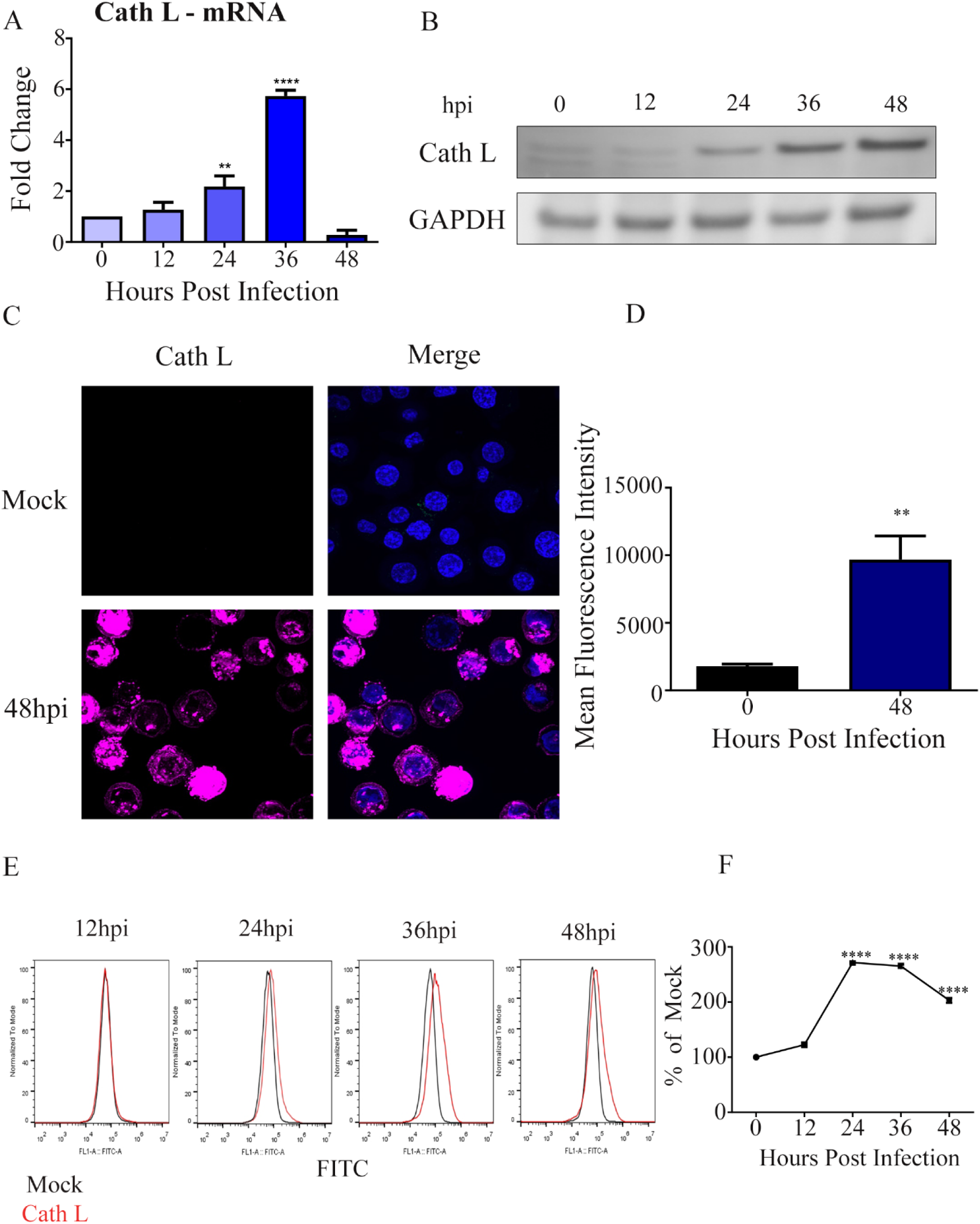
Cathepsin L as mechanism of HPSE activation. **A**. Increase in Cathepsin L mRNA levels, shown is the average fold increase over uninfected control. **B**. Representative western blot showing increase of Cathepsin L over time after infection with HSV-2 333 after a MOI of 1 infection. **C**. Representative immunofluorescence microscopy images of Cathepsin L stain. HSV-2 333 was used to infect cells at an MOI of 1 for 24h. Upper left is Cathepsin L stain only in uninfected sample, upper right is hoescht and Cathepsin L stain merged for uninfected, lower left is Cathepsin L stain only for infected sample at 24hpi, lower right is hoescht and Cathepsin L stain merged for infected sample at 48hpi. **D**. Quantification of total cathepsin L expression from multiple immunofluorescent images. **E**. Representative flow cytometry histogram showing change in Cathepsin L expression with red representing infected samples and black showing uninfected control. **F**. Quantification of Cathepsin L flow cytometry experiments. Asterisks denote a significant difference as determined by Student’s *t*-test; ^∗^*P*<0.05, ^∗∗^*P*<0.01, ^∗∗∗^*P*<0.0001, ^∗∗∗∗^P<0.0001.

### Effect of inhibition of cathepsin L and HPSE on infection

Having established the mechanism through which HPSE is upregulated and activated, we wanted to assess whether the inhibition of HPSE and cathepsin L would affect HSV-2 viral lifecycle. To ascertain the role of HPSE we used a well characterized and commercially available small molecule HPSE activity inhibitor, OGT 2115. This compound functionally blocks HPSE activity, and does not significantly affect its expression (26). To verify the inhibition of HPSE activity during infection we analyzed the cell surface HS expression. While cell surface HS expression during HSV-2 infection usually decreases, our results after pharmacological inhibition of HPSE showed (Figure 5A) a drastic increase in HS expression in both HSV-2 and mock infected cells (Figure 5B). Furthermore, we also observed a decrease in HSV-2 infection as measured by GFP reporter activity in the presence of OGT 2115 at 10 μM concentration. (Figure 5C, 5D). Immunofluorescence microscopy data were in accordance with these results when we used the same HSV-2 GFP reporter virus and the same OGT 2115 concentration. (Figure 5E). We also observed a significant decrease in virus production and release using cell culture supernatant plaque assays through 48 hpi (Figure 5F). These results suggest that pharmacological inhibition of HPSE using OGT 2115 significantly increases HS expression on the cell surface while reducing the overall viral egress and spread.

**Figure 5.**
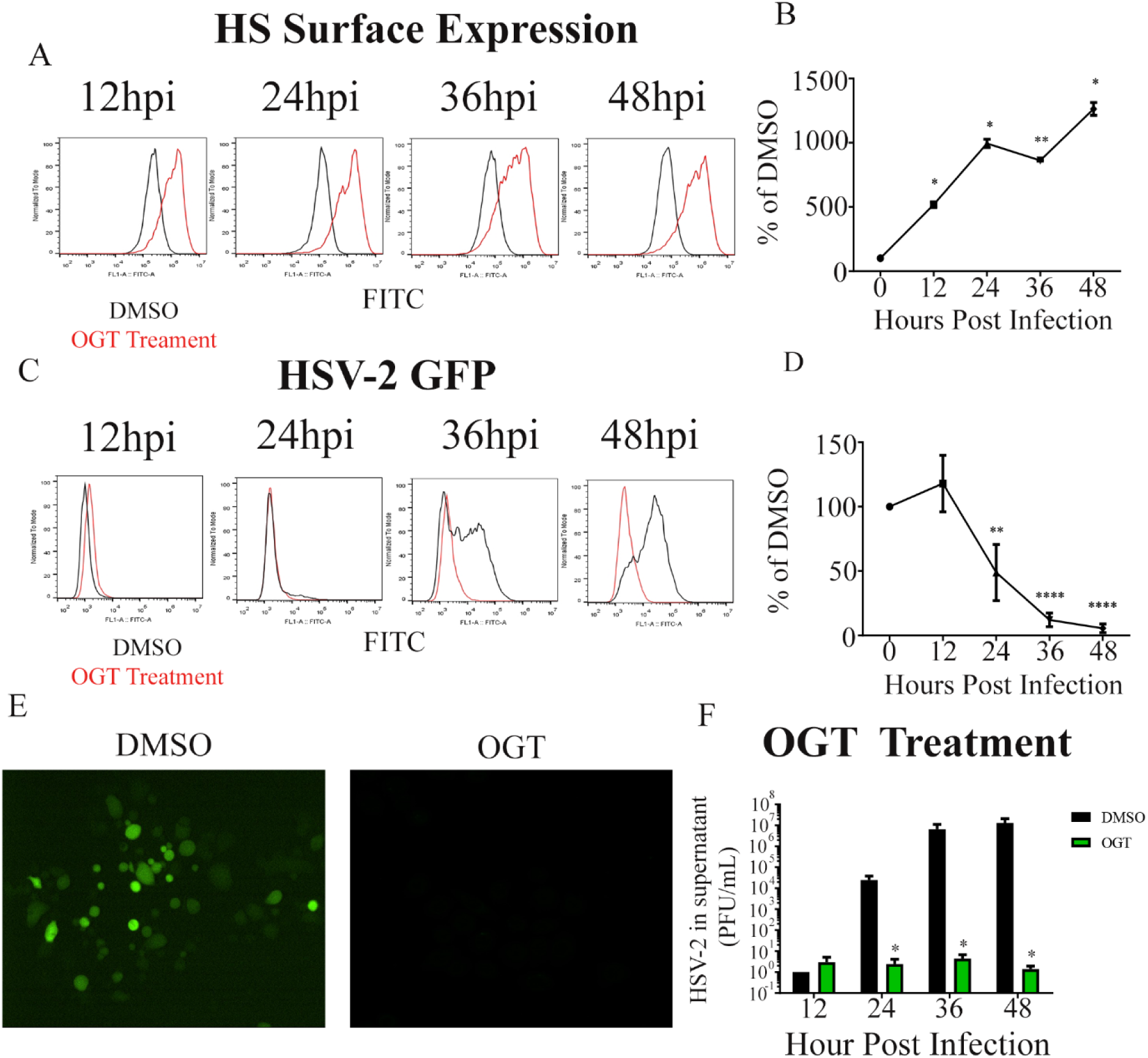
Inhibition of HPSE on HSV-2 infection. **A**. Representative flow cytometry histogram showing change in Cell surface HS expression with red denoting OGT 2115 treatment and Black representing DMSO treatment. Both samples were infected with HSV-2 333 at a MOI of 1. **B**. Quantification of cell surface HS OGT flow cytometry experiments. **C**. Representative flow cytometry histogram showing change in infection following OGT treatment, with red denoting OGT 2115 treatment and Black representing DMSO treatment. Both samples were infected with HSV-2 333 GFP. **D**. Quantification of infection after treatment with OGT flow cytometry experiments. **E**. Representative immunofluorescence microscopy images taken 24hpi after infection with HSV-2 333 GFP after treatment with OGT or DMSO with green representing infected cells. **F**. Average change in virus released from cells with and without OGT treatment every 12h up till 48h. Asterisks denote a significant difference as determined by Student’s *t*-test; ^∗^*P*<0.05, ^∗∗^*P*<0.01, ^∗∗∗^*P*<0.001, ^∗∗∗∗^P<0.0001.

Next we wanted to understand if the inhibition of cathepsin L, the lysosomal activator of HPSE, would affect HSV-2 viral egress and spread. We used a well characterized and commercially available small molecule inhibitor of cathepsin L, cathepsin L inhibitor IV (27). Given that active HPSE remains in the lysosomal compartment for a period of 48 hours before it is degraded (28), we hypothesized that cathepsin L would need to be inhibited for a period up to 48 hours prior to and during infection to make sure no active HPSE is present throughout. As hypothesized, we saw a loss of infection in cathepsin L inhibitor IV treated cells compared to mock DMSO control. These results were consistent when analyzed through immunofluorescence microscopy (Figure 6A) and flow cytometry (Figure 6C, 6D). We also observed through plaque assay that the amount of egressed virus, found in the infected cell supernatant, also decreased in cathepsin L inhibitor IV treated cells compared to mock DMSO treated samples, reaching significance at 48 hpi (Figure 6B).

**Figure 6.**
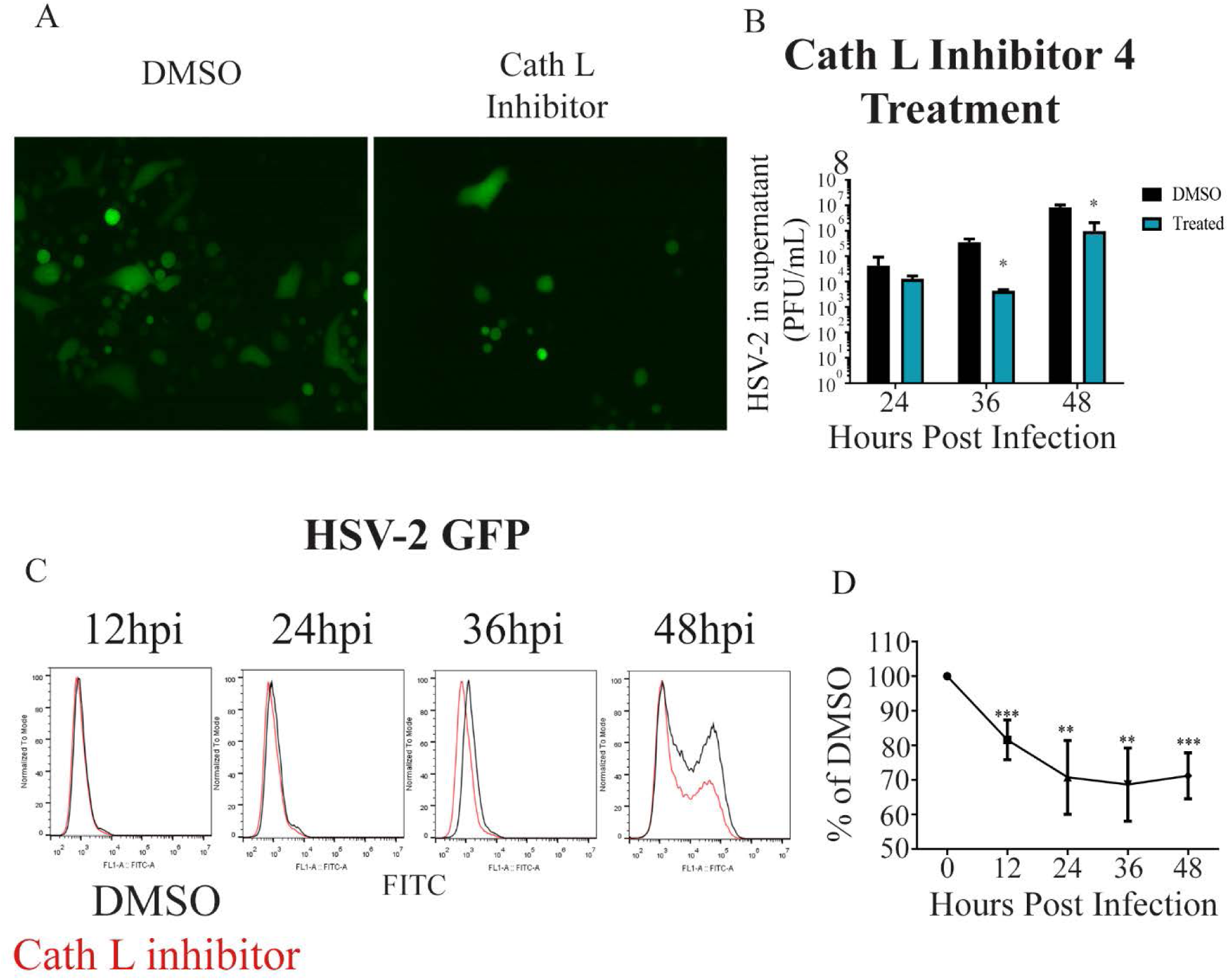
Inhibition of Cathepsin L on HSV-2 infection. **A**. Representative immunofluorescence microscopy images taken 24hpi after infection with HSV-2 333 GFP after treatment with Cathepsin L inhibitor IV or DMSO with green representing infected cells. **B**. Average change in virus released from cells with and without Cathepsin L inhibitor IV treatment every 12h up till 48h. **C**. Representative flow cytometry histogram showing change in infection after Cathepsin L inhibitor IV treatment, with red denoting Cathepsin L inhibitor IV treatment and Black representing DMSO treatment. Both samples were infected with HSV-2 333 GFP. **D**. Quantification of infection after treatment with Cathepsin L inhibitor IV flow cytometry experiments. Asterisks denote a significant difference as determined by Student’s *t*-test; ^∗^*P*<0.05, ^∗∗^*P*<0.01, ^∗∗∗^*P*<0.001.

### Effect of overexpression of HPSE during HSV-2 infection

Through the experiments in previous sections, we were able to establish that HPSE is important for HSV-2 release. Lastly, we wanted to understand if the overexpression of HPSE during a HSV-2 infection would be beneficial during viral proliferation. To study this, we overexpressed HPSE in VK2 cells for a period of 24 hours followed by infection with HSV-2 (333) GFP virus at a MOI of 1. The cells were plated without any methylcellulose to allow release and spread of the extracellular virions. We observed via immunofluorescence microscopy a clear increase in virally infected cell clusters in the HPSE transfected cells when compared to empty vector control (Figure 7A). It is important to note that although we saw less infection than expected, we normally find less infection in samples that have been previously transfected. Our results were in accordance with flow cytometry results (Figure 7C), which showed that overexpression of HPSE lead to a faster spread of infection, reaching significance at 36 and 48 hpi (Figure 7D). We also observed an increase in released virus after HPSE overexpression (Figure 7B) at 36 hpi and 48 hpi. Taken together our results suggest a direct connection between higher levels of HPSE and HSV-2 release in the culture supernatant.

**Figure 7.**
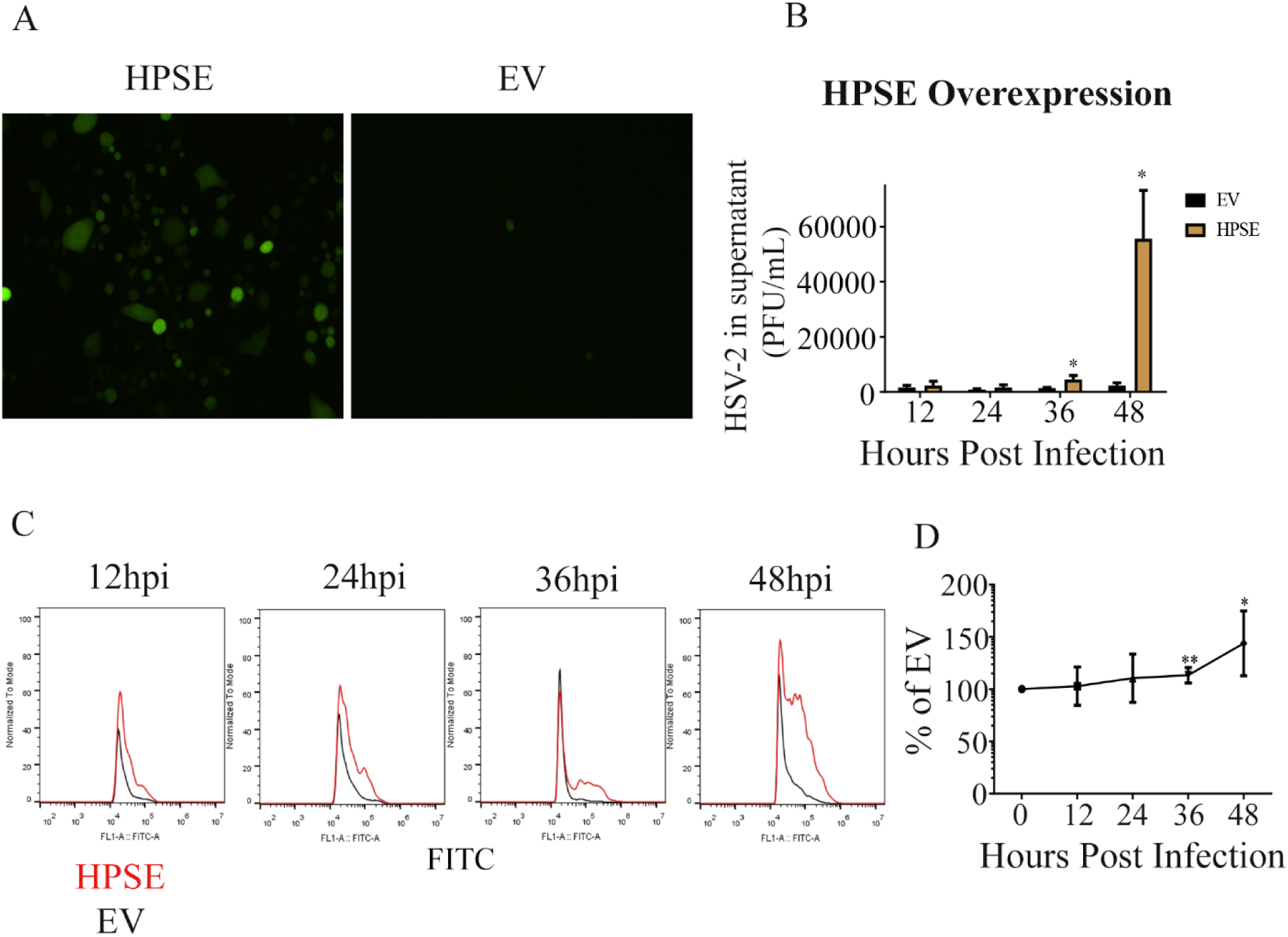
Effect of overexpression of HPSE during HSV-2 infection. **A**. Representative immunofluorescence microscopy images taken 24hpi after infection with HSV-2 333 GFP after overexpression of HPSE with green representing infected cells. **B**. Average change in virus released from cells overexpressing HPSE then infected with HSV-2 333 every 12h up till 48h. **C**. Representative flow cytometry histogram showing change in newly infected cells after overexpression of HPSE, with red denoting HPSE overexpression and black representing EV. Both samples were infected with HSV-2 333 GFP. **D**. Quantification of infection after overexpression of HPSE flow cytometry experiments. Asterisks denote a significant difference as determined by Student’s *t*-test; ^∗^*P*<0.05, ^∗∗^*P*<0.01.

## Discussion

Active HPSE is responsible for the degradation of cell surface HS. HS is found covalently attached to a small set of extracellular matrix and plasma membrane proteins forming heparan sulfate proteoglycans (HSPG) (15, 16). Clearance of HS via HPSE modulates cell division and differentiation, tissue morphogenesis and architecture, and organismal physiology (16). HSV-2 encodes for two envelope glycoproteins, gB and gC, which bind HS at the cell surface and initiate viral entry (17-19). However, the new virions released can face a significant challenge if they bind with HS during viral egress. This phenomenon is commonly seen with other viruses including influenza where the viral hemagglutinin binds to host sialic acid chains and restricts viral release (32, 33). To eliminate the possibility of new virions binding upon release, the surface sugars are cleaved by a viral endoglycosidase (34, 35). In this regard, HSV-2 does not have any known viral proteins that can cleave cell surface HS. HSV-2 may have found the ability to cleave HS nevertheless from a host enzyme (20, 21). Given that HPSE is the only known mammalian enzyme capable of cleaving HS chains (9), we hypothesized and then demonstrated that its upregulation during infection may be beneficial for HSV-2 release from host cells.

Our evidence of HPSE upregulation during infection may suggest an additional role for HPSE in the exacerbation of the genital disease caused by HSV-2. During genital infection, distressing, painful lesions, and sores around the genitals or rectum can result from a productive HSV-2 infection. These lesions are accompanied by inflammation and localized loss to tissue architecture (6). Recent studies have implicated HPSE overexpression in multiple pathologic processes, including inflammation (36), angiogenesis (37), tumor metastasis (38), and atherosclerosis (39). It has been shown how heparanase contributes to HS cleavage and how this specifically results in rearrangement of the extracellular matrix as well as in controlling the release of many cell surface HS-linked molecules such as growth factors, cytokines and enzymes in larger tissue wide changes (9, 40).

To understand potential contributions of HPSE during HSV-2 infection, we first looked at HS chain expression on the cell surface during HSV-2 infection. It became very clear to us that HS expression decreases over time during infection. This information also strengthened our goal to see how HPSE is modulated during infection. As suspected, HPSE was upregulated both transcriptionally and translationally showing higher protein levels on the cell surface at 24, 36 and 48 hpi. Together, these results suggest that HSV-2 infected cells upregulate HPSE, which is then translocated to the cell surface, decreasing HS on the cell surface after infection.

Next we wanted to understand the mechanism behind the upregulation and activation of HPSE during infection. It was reported that nuclear factor NF-κB, is upregulated upon HSV infection and that NF-κB upregulation has been linked to the transcriptional regulation of HPSE (20). Also, it is well known that cathepsin L, a lysosomal endopeptidase, is the only known activator of HPSE in mammalian cells. Hence we looked at the transcriptional and protein levels of these two factors and how they change with infection. Through this study, we were able to show that HSV-2 infection of VK2 cells upregulated NF-κB and cathepsin L which in turn orchestrated the expression and activation of HPSE.

Finally, we asked whether the inhibition of HPSE and cathepsin L would downregulate viral egress and spread. To study this, we used two well-known inhibitors, OGT 2115 and cathepsin L inhibitor IV, on VK2 cells during HSV-2 infection. We saw a significant loss of infection using both these inhibitors compared to vehicle controls suggesting an alternate therapeutic modality against HSV-2 infections. Additionally, when we upregulated HPSE expression using a HPSE expression plasmid, we saw a significant increase in viral progeny and spread.

For our model we propose that during the productive phase of HSV-2 infection of VK2 cells HSV-2 causes an upregulation in HPSE levels that is mediated by an increased NF-κB p65 nuclear localization and a resultant activation of HPSE transcription. In parallel, increased levels of cathepsin L contribute to HPSE proenzyme cleavage and activation. Together they contribute to the removal of heparan sulfate from cell surface and thus, facilitate virus release from cells. Higher levels of HPSE may also be a trigger for the breakdown of extracellular matrix and eventually a trigger for local inflammation. While the latter part is yet to be demonstrated, our studies discussed here directly implicate HPSE and cathepsin L in HSV-2 release.

## Materials and Methods

### Cells and viruses

Human vaginal epithelial (VK2/E6E7) cells obtained from ATCC. VK2/E6E7 cells were passaged in Karatinocyte serum free medium (KSFM) (Gibco/BRL, Carlsbad, CA, USA) supplemented with epidermal growth factor (EGF), bovine pituitary extract (BPE) and 1% penicillin/streptomycin. For convenience this cell line is referred to as VK2 cells throughout. All infections were done with HSV-2 333 at MOI of 0.1 or 1 on VK2 cells unless mentioned otherwise. The Vero cell line cell line (African green monkey kidney) was generously given by Dr. Patricia G. Spear (Northwestern University, Chicago, IL) and cultured in DMEM (Gibco) with 10% FBS and 1% penicillin/streptomycin. Cathepsin L inhibitor IV (Santa Cruz Biotechnology) was used as a cathepsin L activity inhibitor (27). Cathepsin L inhibitor was used at a concentration of 10 μg/mL in DMSO unless otherwise specified. OGT 2115 (Tocris Biosciences) was used for heparanase activity inhibition and has been previously described as a HPSE inhibitor (<u>41</u>). OGT 2115 was used at 10 μM unless otherwise specified. Viruses used were wild type HSV-2 (333), and HSV-2 (333) GFP (42). Virus stocks were grown and tittered on Vero cells, and stored at –80°C.

### Antibodies and plasmids

HPSE antibody H-80 (Santa Cruz Biotechnology) was used for imaging (1:100) and flow cytometry studies (1:100). Cathepsin L antibody ab58991 (Abcam, Cambridge MA) was used for western blot analysis (1:2,000), imaging (1:100) and flow cytometry studies (1:200). Histone H3 antibody (Cell Signaling Technology, Danvers, MA) was used at a dilution of 1:500 for western blot analysis. Anti-human HS monoclonal antibody 10E4 (US Biological, Salem, MA) was used for flow cytometry (1:100) and cell imaging (1:100). NF-κB p65 antibody C-20 (Santa Cruz Biotechnology) was used for imaging (1:500) and western blot analysis (1:2,000). GAPDH (Santa Cruz Biotechnology) was used for western blot analysis at dilution 1:2,000. The IkBa (S32/36A) plasmid was provided by Dr Michael Karin (University of California, San Diego, La Jolla, CA). All transfections were performed using Lipofectamine-2000 (Life Technologies).

### Western blot analysis

Proteins from samples in this study were collected using radio immunoprecipitation assay (RIPA) buffer (Sigma-Aldrich, St Louis, MO) according to the manufacturer’s protocol. After gel electrophoresis, membranes were blocked in 5% BSA for 1 h followed by incubation with primary antibody for 1 h, then incubation with respective secondary antibodies (anti-mouse 1:10,000, anti-rabbit 1:10,000) for 1 h. Protein bands were visualized using the SuperSignal West Femto maximum sensitivity substrate (Thermo Scientific, Waltham, MA) with Image-Quant LAS 4000 biomolecular imager (GE Healthcare Life Sciences, Pittsburgh, PA). The densities of the bands were quantified using ImageJ 1.52a image analysis software (NIH, USA).

### PCR

Trizol (Life Technologies) was used to obtain RNA following the protocol from the manufacturer then a cDNA Reverse Transcription Kit from (Applied Biosystems) was used to transcribe to DNA following the protocol from the manufacturer. Fast SYBR Green Master Mix (Applied Biosystems) was used to perform real-time quantitative PCR, using QuantStudio 7 Flex (Applied Biosystems). The primers used in this study are as follows: HPSE forward primer 5’-CTCGAAGAAAGACGGCTA-3’ and reverse primer 5’-GTAGCAGTCCGTCCATTC-3’; NF-κB p65 forward primer 5’-TGGGGACTACGACCTGAATG-3’ and reverse primer 5’-GGGGGCACGATTGTCAAAGA-3’; GAPDH forward primer 5’-TCCACTGGCGTCTTCACC-3’ and reverse primer 5’-GGCAGAGATGATGACCCTTTT-3’; cathepsin L forward primer 5’-ATCTGGGCATGAACCACCTG-3’ and reverse primer 5’-CAGCAAGCACCACAAGAACC-3’.

### Luciferase assay

An empty vector, pGL3-Basic (Promega, Madison, WI), was used to control for transfection efficiency. The pHep1(0.7 kb)-luc plasmid that expresses firefly luciferase that is driven by the 0.7-kb upstream promoter for transcription of human HPSE was provided by Dr Xiulong Xu (Rush University). Luciferase activity was measured using Berthold detection systems Luminometer.

### Flow cytometry

Measurement of HS and HPSE cell surface expression was performed after HSV-2 333-WT infection. Monolayers of VK2 cells were infected at MOI 0.1 or 1 then harvested at different times post infection (12 hpi, 24 hpi, 36 hpi, and 48 hpi). Cells were harvested and fixed with 4% PFA for 10 min, then incubated with 5% BSA for 1 h followed by incubation with FITC-conjugated anti-HS diluted 1:100 in PBS with 1% BSA for 1 h. Cells were then suspended in FACS buffer (PBS, 5% FBS, 0.1% sodium azide). For detection of HPSE on the cell surface, cells were harvested then fixed with 4% PFA for 4 min, then incubated with 5% BSA for 1 h, then incubated with primary antibody diluted in PBS with 5% BSA for 1 h, followed by incubation with FITC-conjugated secondary antibody. Cells were then resuspended in FACS buffer. Cells stained with respective FITC-conjugated secondaries only were used as background controls. Entire cell populations were used for the mean fluorescence intensity calculations.

### Immunofluorescence microscopy

VK2 cells were cultured in glass bottom dishes (MatTek Corporation, Ashland, MA). Cells were fixed in 4% PFA for 10 min and permeabilized with 0.1% Triton-X for intracellular labelling, then incubated with 5% BSA for 1 hour then primary antibody diluted in PBS with 5% BSA for 1 hour then with FITC-conjugated secondary. Then imaging was done with a Zeiss Confocal 710, Germany. Pinhole was set to 1 Airy Unit. Fluorescence intensity of images was calculated using Zen software.

### Plaque assay

Viral egress titers were measured using a plaque assay. Monolayers of VK2 cells were plated in twelve-well plates and infected with HSV-2 333 virus at MOI 0.1 or 1. Media were collected at different time points post infection and titered on Vero cells. Primary incubation of collected media was performed with DPBS+/+ (Life Technologies) with 1% glucose and 1% heat-inactivated serum for 2 h. Vero cells were then incubated with growth media containing 5% methylcellulose for 72 h followed by fixing with 100% methanol and staining with crystal violet solution.

### Statistics

Error bars of all figures represent standard error of three independent experiments, unless otherwise specified. Asterisks denote a significant difference as determined by Student’s *t*-test; ^∗^P<0.05, ^∗∗^P<0.01, ^∗∗∗^P<0.001, ^∗∗∗∗^p<0.0001, ns, not significant.

## Acknowledgments

The authors acknowledge Ruth Zhelka for help with using the departmental imaging facilities. This work was supported by a grant from the NIH (R01 1R21AI128171-02) to D.S. J.H. was supported by ISPB grant G3108. T.Y. was supported by ISPB grant G3126. A.A. was supported by fellowship F30EY025981.

## References

1. F Xu, M., S. MR, G. SL, B. SM, M. LE, F. SE, and T. LD. 2010. Seroprevalence of Herpes Simplex Virus Type 2 Among Persons Aged 14–49 Years — United States, 2005–2008. Morbidity and Mortality Weekly Report (MMWR). 59:456–457,458,459.

2. Kinghorn, G. R. 1993. Genital herpes: natural history and treatment of acute episodes. J. Med. Virol. Suppl 1:33–38.

3. Scoular, A., J. Norrie, G. Gillespie, N. Mir, and W. F. Carman. 2002. Longitudinal study of genital infection by herpes simplex virus type 1 in Western Scotland over 15 years. BMJ. 324:1366–1367.

4. Wald, A. 2006. Genital HSV-1 infections. Sex. Transm. Infect. 82:189–190. doi:82/3/189.

5. Xu, F., M. R. Sternberg, B. J. Kottiri, G. M. McQuillan, F. K. Lee, A. J. Nahmias, S. M. Berman, and L. E. Markowitz. 2006. Trends in herpes simplex virus type 1 and type 2 seroprevalence in the United States. JAMA. 296:964–973. doi:296/8/964.

6. Jaishankar, D., and D. Shukla. 2016. Genital Herpes: Insights into Sexually Transmitted Infectious Disease. Microb. Cell. 3:438–450. doi: 10.15698/mic2016.09.528 [doi].

7. Halpern-Felsher, B. L., J. L. Cornell, R. Y. Kropp, and J. M. Tschann. 2005. Oral versus vaginal sex among adolescents: perceptions, attitudes, and behavior. Pediatrics. 115:845–851. doi: 115/4/845 [pii].

8. Strick, L. B., A. Wald, and C. Celum. 2006. Management of herpes simplex virus type 2 infection in HIV type 1-infected persons. Clin. Infect. Dis. 43:347–356. doi: CID39058.

9. Vlodavsky, I., N. Ilan, A. Naggi, and B. Casu. 2007. Heparanase: structure, biological functions, and inhibition by heparin-derived mimetics of heparan sulfate. Curr. Pharm. Des. 13:2057–2073.

10. Fairbanks, M. B., A. M. Mildner, J. W. Leone, G. S. Cavey, W. R. Mathews, R. F. Drong, J. L. Slightom, M. J. Bienkowski, C. W. Smith, C. A. Bannow, and R. L. Heinrikson. 1999. Processing of the human heparanase precursor and evidence that the active enzyme is a heterodimer. J. Biol. Chem. 274:29587–29590.

11. Wu, L., C. M. Viola, A. M. Brzozowski, and G. J. Davies. 2015. Structural characterization of human heparanase reveals insights into substrate recognition. Nat. Struct. Mol. Biol. 22:1016–1022. doi: 10.1038/nsmb.3136.

12. Fux, L., N. Feibish, V. Cohen-Kaplan, S. Gingis-Velitski, S. Feld, C. Geffen, I. Vlodavsky, and N. Ilan. 2009. Structure-function approach identifies a C-terminal domain that mediates heparanase signaling. Cancer Res. 69:1758–1767. doi: 10.1158/0008-5472.CAN-08-1837.

13. Coulombe, R., P. Grochulski, J. Sivaraman, R. MÃ©nard, J. S. Mort, and M. Cygler. 1996. Structure of human procathepsin L reveals the molecular basis of inhibition by the prosegment. EMBO J. 15:5492–5503.

14. Ishidoh, K., T. Towatari, S. Imajoh, H. Kawasaki, E. Kominami, N. Katunuma, and K. Suzuki. 1987. Molecular cloning and sequencing of cDNA for rat cathepsin L. FEBS Lett. 223:69–73. doi: 0014-5793(87)80511-2.

15. Bishop, J. R., M. Schuksz, and J. D. Esko. 2007. Heparan sulphate proteoglycans fine-tune mammalian physiology. Nature. 446:1030–1037. doi: nature05817.

16. Esko, J. D., and S. B. Selleck. 2002. Order out of chaos: assembly of ligand binding sites in heparan sulfate. Annu. Rev. Biochem. 71:435–471. doi: 10.1146/annurev.biochem.71.110601.135458.

17. Spear, P. G., R. J. Eisenberg, and G. H. Cohen. 2000. Three classes of cell surface receptors for alphaherpesvirus entry. Virology. 275:1–8. doi: 10.1006/viro.2000.0529.

18. Gruenheid, S., L. Gatzke, H. Meadows, and F. Tufaro. 1993. Herpes simplex virus infection and propagation in a mouse L cell mutant lacking heparan sulfate proteoglycans. J. Virol. 67:93–100.

19. Spear, P. G., M. T. Shieh, B. C. Herold, D. WuDunn, and T. I. Koshy. 1992. Heparan sulfate glycosaminoglycans as primary cell surface receptors for herpes simplex virus. Adv. Exp. Med. Biol. 313:341–353.

20. Hadigal, S. R., A. M. Agelidis, G. A. Karasneh, T. E. Antoine, A. M. Yakoub, V. C. Ramani, A. R. Djalilian, R. D. Sanderson, and D. Shukla. 2015. Heparanase is a Host Enzyme Required for Herpes Simplex Virus-1 Release from Cells. Nat. Commun. 6:6985. doi: 10.1038/ncomms7985.

21. Guo, C., Z. Zhu, Y. Guo, X. Wang, P. Yu, S. Xiao, Y. Chen, Y. Cao, and X. Liu. 2017. Heparanase upregulation contributes to porcine reproductive and respiratory syndrome virus release. J. Virol. doi: JVI.00625-17.

22. Kasperczyk, H., K. La Ferla-Bruhl, M. A. Westhoff, L. Behrend, R. M. Zwacka, K. M. Debatin, and S. Fulda. 2005. Betulinic acid as new activator of NF-kappaB: molecular mechanisms and implications for cancer therapy. Oncogene. 24:6945–6956. doi: 1208842.

23. Goldshmidt, O., E. Zcharia, R. Abramovitch, S. Metzger, H. Aingorn, Y. Friedmann, V. Schirrmacher, E. Mitrani, and I. Vlodavsky. 2002. Cell surface expression and secretion of heparanase markedly promote tumor angiogenesis and metastasis. Proc. Natl. Acad. Sci. U. S. A. 99:10031–10036. doi: 152070599.

24. DiDonato, J. A., M. Hayakawa, D. M. Rothwarf, E. Zandi, and M. Karin. 1997. A cytokine-responsive IkappaB kinase that activates the transcription factor NF-kappaB. Nature. 388:548–554. doi: 10.1038/41493.

25. Bakhshi, R., A. Goel, P. Seth, P. Chhikara, and S. S. Chauhan. 2001. Cloning and characterization of human cathepsin L promoter. Gene. 275:93–101. doi: http://dx.doi.org/10.1016/S0378-1119(01)00650-3.

26. Courtney, S. M., P. A. Hay, R. T. Buck, C. S. Colville, D. J. Phillips, D. I. Scopes, F. C. Pollard, M. J. Page, J. M. Bennett, M. L. Hircock, E. A. McKenzie, M. Bhaman, R. Felix, C. R. Stubberfield, and P. R. Turner. 2005. Furanyl-1,3-thiazol-2-yl and benzoxazol-5-yl acetic acid derivatives: novel classes of heparanase inhibitor. Bioorg. Med. Chem. Lett. 15:2295–2299. doi: S0960-894X(05)00295-7.

27. Yasuma, T., S. Oi, N. Choh, T. Nomura, N. Furuyama, A. Nishimura, Y. Fujisawa, and T. Sohda. 1998. Synthesis of peptide aldehyde derivatives as selective inhibitors of human cathepsin L and their inhibitory effect on bone resorption. J. Med. Chem. 41:4301–4308. doi: 10.1021/jm9803065.

28. Abboud-Jarrous, G., R. Atzmon, T. Peretz, C. Palermo, B. B. Gadea, J. A. Joyce, and I. Vlodavsky. 2008. Cathepsin L Is Responsible for Processing and Activation of Proheparanase through Multiple Cleavages of a Linker Segment. J. Biol. Chem. 283:18167–18176. doi: 18167.

29. Shukla, D., and P. G. Spear. 2001. Herpesviruses and heparan sulfate: an intimate relationship in aid of viral entry. J. Clin. Invest. 108:503–510. doi: 10.1172/JCI13799.

30. Weissenhorn, W., A. Hinz, and Y. Gaudin. 2007. Virus membrane fusion. FEBS Lett. 581:2150–2155. doi: S0014-5793(07)00166-4.

31. Agelidis, A. M., and D. Shukla. 2015. Cell entry mechanisms of HSV: what we have learned in recent years. Future Virol. 10:1145–1154. doi: 10.2217/fvl.15.85.

32. Suzuki, Y., T. Ito, T. Suzuki, R. E. Holland, T. M. Chambers, M. Kiso, H. Ishida, and Y. Kawaoka. 2000. Sialic Acid Species as a Determinant of the Host Range of Influenza A Viruses. J. Virol. 74:11825–11831. doi: 0798.

33. Chen, L. M., O. Blixt, J. Stevens, A. S. Lipatov, C. T. Davis, B. E. Collins, N. J. Cox, J. C. Paulson, and R. O. Donis. 2012. In vitro evolution of H5N1 avian influenza virus toward human-type receptor specificity. Virology. 422:105–113. doi: 10.1016/j.virol.2011.10.006.

34. Baum, L. G., and J. C. Paulson. 1991. The N2 neuraminidase of human influenza virus has acquired a substrate specificity complementary to the hemagglutinin receptor specificity. Virology. 180:10–15.

35. Schulman, J. L. 1969. The role of antineuraminidase antibody in immunity to influenza virus infection. Bull. World Health Organ. 41:647–650.

36. Changyaleket, B., Z. Z. Chong, R. O. Dull, D. Nanegrungsunk, and H. Xu. 2017. Heparanase promotes neuroinflammatory response during subarachnoid hemorrhage in rats. J. Neuroinflammation. 14:137-017-0912-8. doi: 10.1186/s12974-017-0912-8.

37. Poupard, N., P. Badarou, F. Fasani, H. Groult, N. Bridiau, F. Sannier, S. Bordenave-Juchereau, C. Kieda, J. M. Piot, C. Grillon, I. Fruitier-Arnaudin, and T. Maugard. 2017. Assessment of Heparanase-Mediated Angiogenesis Using Microvascular Endothelial Cells: Identification of lambda-Carrageenan Derivative as a Potent Anti Angiogenic Agent. Mar. Drugs. 15:10.3390/md15050134. doi: E134.

38. Jiao, F., S. Bai, Y. Ma, Z. Yan, Z. Yue, Y. Yu, X. Wang, and J. Wang. 2014. DNA Methylation of Heparanase Promoter Influences Its Expression and Associated with the Progression of Human Breast Cancer. PLoS One. 9:e92190. doi:10.1371/journal.pone.0092190. doi: PONE-D-13-40068.

39. Planer, D., S. Metzger, E. Zcharia, I. D. Wexler, I. Vlodavsky, and T. Chajek-Shaul. 2011. Role of heparanase on hepatic uptake of intestinal derived lipoprotein and fatty streak formation in mice. PLoS One. 6:e18370. doi: 10.1371/journal.pone.0018370.

40. Sanderson, R. D., M. Elkin, A. C. Rapraeger, N. Ilan, and I. Vlodavsky. 2017. Heparanase regulation of cancer, autophagy and inflammation: new mechanisms and targets for therapy. The FEBS Journal. 284:42–55. doi: 10.1111/febs.13932.

41. Courtney, S. M., P. A. Hay, R. T. Buck, C. S. Colville, D. J. Phillips, D. I. C. Scopes, F. C. Pollard, M. J. Page, J. M. Bennett, M. L. Hircock, E. A. McKenzie, M. Bhaman, R. Felix, C. R. Stubberfield, and P. R. Turner. 2005. Furanyl-1,3-thiazol-2-yl and benzoxazol-5-yl acetic acid derivatives: novel classes of heparanase inhibitor. Bioorganic & Medicinal Chemistry Letters. 15:2295–2299. doi: 10.1016/j.bmcl.2005.03.014. http://www.sciencedirect.com/science/article/pii/S0960894X05002957.

42. Wanda M. Martinez, and Patricia G. Spear. 2002. Amino Acid Substitutions in the V Domain of Nectin-1 (HveC) That Impair Entry Activity for Herpes Simplex Virus Types 1 and 2 but Not for Pseudorabies Virus or Bovine Herpesvirus 1. Journal of Virology. 76:7255–7262. doi: 10.1128/JVI.76.14.7255-7262.2002. http://jvi.asm.org/content/76/14/7255.abstract.

